# Beyond the nucleus: Plastic chemicals activate G protein-coupled receptors

**DOI:** 10.1101/2023.10.04.560665

**Authors:** Molly McPartland, Sarah Stevens, Zdenka Bartosova, Ingrid Gisnås Vardeberg, Johannes Völker, Martin Wagner

**Author notes:** Corresponding author: Molly McPartland, Martin Wagner.

## Abstract

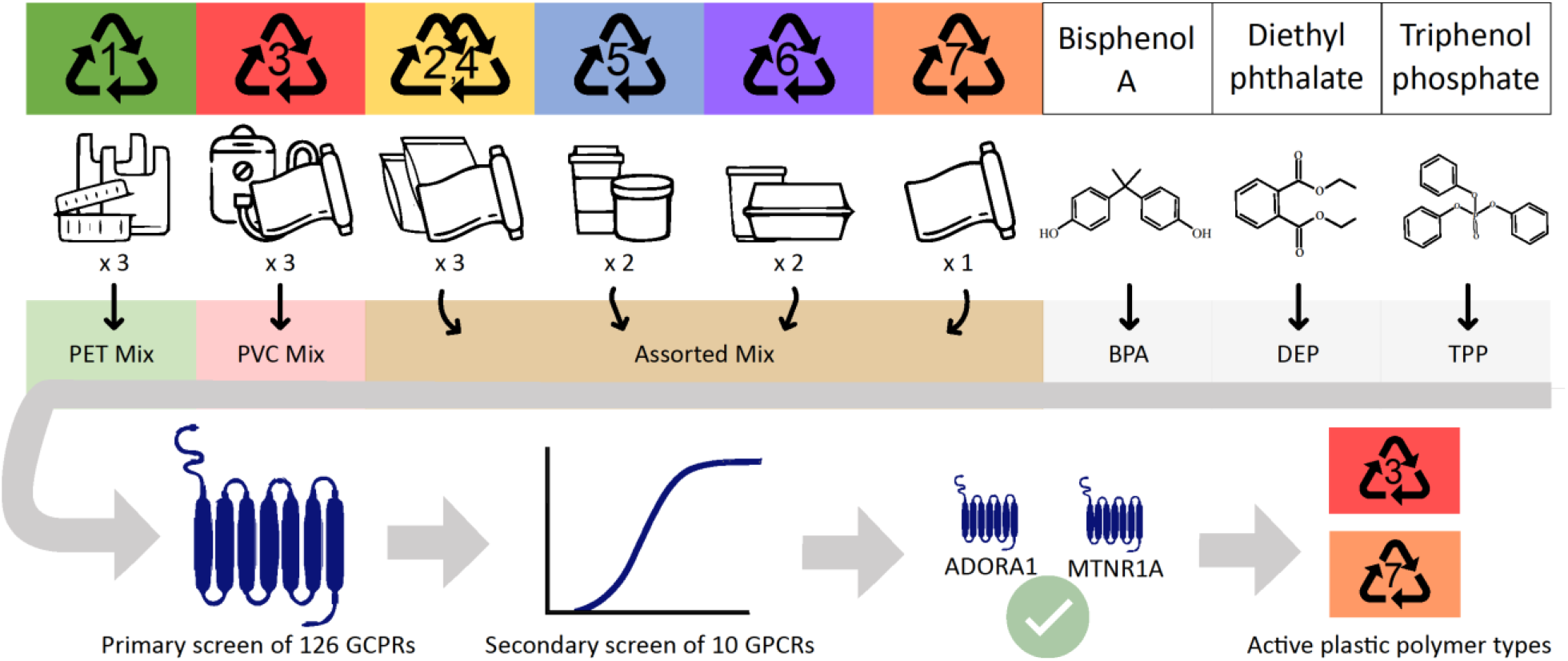

G protein-coupled receptors (GPCRs) are central mediators of cell signaling and physiological function. Despite their biological significance, GPCRs have not been widely studied in the field of toxicology. Herein, we investigated these receptors are novel targets of plastic chemicals using a high-throughput drug screening assay with 126 human non-olfactory GPCRs. In a first-pass screen, we tested the activity of triphenol phosphate, bisphenol A, and diethyl phthalate as well as three real-world mixtures of chemicals extracted from plastic food packaging covering all major polymer types. We found 11 GPCR-chemical interactions, of which the chemical mixtures exhibited the most robust activity at Adenosine receptor 1 (ADORA1) and Melatonin receptor 1 (MTNR1A) in a confirmatory secondary screen. We further confirmed that polyvinyl chloride and polyurethane products contain ADORA1 or MTNRA1 agonists using pharmacological knockdown experiments. Finally, an analysis of the associated gene ontology terms suggests that ADORA1 and MTNR1A activation may be linked to downstream effects on circadian and metabolic processes. Our findings exemplify the diversity of endpoints plastic chemicals can target and demonstrate the relevance of non-genomic pathways which have thus far remained unexplored.

## 1 Introduction

G protein-coupled receptors (GPCRs) are the largest class of cell surface receptors and transduce signals from a diverse array of ligands across the cell membrane. They function as central regulators for most cellular processes, including cell differentiation, migration, apoptosis, and growth.^1^ Physiologically, disruption of GPCR signaling is linked to many diseases, underscored by the fact that one-third of all pharmaceuticals target GPCRs.^2^ Despite their biological significance, GPCRs have received very little attention in toxicological research. This is striking given that endocrine disrupting chemicals (EDCs), compounds that can “interfere with any aspect of hormone action”,^3^ often act via cellular receptors, namely nuclear receptors, and represent a major topic of research.

Indeed, emerging evidence suggests that EDCs or synthetic chemicals can interact with several rhodopsin-like GPCRs as well. A prominent example is Bisphenol A (BPA) activating G protein-coupled estrogen receptor 1 (GPER) leading to various cancer-promoting mechanisms, such as increased reactive oxygen species,^4^ cellular proliferation,^5^ apoptosis,^6^ and cell migration.^7^ Beyond GPER and bisphenols, scattered evidence suggests that other chemicals can also interfere with GPCR signaling. For instance, p,p’-DDT allosterically activates the human follitropin receptor (FSHR),^8^ multiple phthalate esters inhibit the cannabinoid-1 (CB_1_) receptor,^9^ and certain carbamate insecticides activate or inhibit melatonin receptors.^10,11^ While not comprehensive, these reports demonstrate that GPCRs are susceptible to known EDCs and other synthetic chemicals.

Plastics are a common source of human exposure to chemicals, including EDCs.^12,13,14^ Well-studied plastic chemicals, such as BPA, have been detected in over 90% of US, European, and Asian populations.^15^ Such exposure is linked to an increased prevalence of non-communicable diseases including asthma, obesity and diabetes, hormone-sensitive carcinogenesis, and impaired immune function.^16^ The associated health costs of exposure to EDCs are estimated at 340 billion US dollars and 160 billion Euros, annually.^17^ This shows that certain plastic chemicals significantly contribute to the burden of disease.

Plastics can contain thousands of chemicals in addition to phthalates and BPA,^18^ including both intentionally-added substances, such as plasticizers, colorants, stabilizers, and flame retardants,^19^ and non-intentionally added substances (NIAS), such as unreacted monomers, reaction byproducts, degradation products, and impurities.^20^ As these chemicals are not covalently bound to the polymer, they can leach into liquids, solids, or air via migration or volatilization, resulting in human exposure to both known and unknown chemicals. Thus, realistic exposure scenarios must consider complex mixtures of plastic chemicals. Assessing toxicity of the overall mixture of compounds released by plastics encompasses all exposure-relevant chemicals and is key to understand their joint impacts and hazard.^21^

In contrast to the dedicated research on plastic chemical mediating their toxicity via nuclear receptors, an interrogation of such chemicals across human GPCRs has not yet been undertaken. Given there are ∼400 non-olfactory GPCRs that may, in principle, be targeted by plastic chemicals, a high-throughput screening of many GPCRs represents an ideal approach. To this end, we adapted the PRESTO-Tango (Parallel Receptor-ome Expression and Screening via Transcriptional Output, with transcriptional activation following beta arrestin translocation) platform used in drug development^22^ to investigate whether plastic chemicals activate 126 non-olfactory GPCRs. To address the chemical complexity of plastic products, including unknown substances and potential mixture toxicity, we evaluated the GPCR agonism caused by all extractable chemicals in plastic food contact articles (FCA) in addition to individual well-known chemicals used in plastics.

In this work, we aimed to investigate (i) whether chemicals present in plastic FCA activate specific GPCRs, (ii) whether certain polymer types or products contain such GPCR agonists, and (iii) the potential biological implications of GPCR activation. Here, we identify several novel receptor-chemical interactions which were confirmed via dose-dependent activation and pharmacological knock-down using known GPCR antagonists. We further identify specific polymer types and products containing potent GPCR agonists. Finally, we identify biological processes associated with the respective GPCR disruption.

## 2 Materials and methods

### 2.1 Rationale and study design

In contrast to the conventional method of testing multiple samples on a single target, we utilized the PRESTO-Tango platform to assess many GPCRs using a smaller set of samples. Accordingly, we screened three individual plastic chemicals and three chemical Mixes derived from extracts of plastic FCAs. This approach allowed us to screen many GPCRs in the first round and then, in further iterations with fewer receptors, confirm these hits and determine specific products and polymer types containing GPCR agonists.

We selected BPA (CAS: 80-05-7, 99.5%, Sigma Aldrich), diethyl phthalate (DEP, 84-66-2, 99.5%, Sigma Aldrich), and triphenol phosphate (TPP, 115-86-6, >99%, Sigma Aldrich) because they were (1) present in plastic FCAs, (2) classified as EDCs,^13,18^ and (3) detected in more than 50% of the examined human populations.^23^ To address “real-world” chemicals present in plastic FCAs, we extracted everyday plastic food packaging using methanol (see 2.2). Specifically, we selected the plastic types with the highest global production volumes^24^ (polypropylene (PP), polyethylene (PE), polyethylene terephthalate (PET), polystyrene (PS), polyvinyl chloride (PVC), polyurethane (PUR)) from the four countries with the highest volume of plastic waste per capita (USA, Germany, England, South Korea),^25^ as well as from local grocery stores and suppliers in Norway. We pooled three products made of PET to make the PET Mix and three products made of polyvinyl chloride (PVC Mix). We focused on these polymer types because of their weak and strong respective toxicity at a range of endpoints.^26^ The third Mix (Assorted Mix) contained three PE products, two PP products, two PS products, and one PUR product (Table 1, Figure S1). With this mixture design, we optimized the number of plastic products we could test and enabled comparison between and within single plastic chemicals and the real-world chemical mixtures present in plastic products.

**Table 1.** Composition of the Mixes of plastic extracts screened in this study and results from the nontarget chemical analysis (number of chemical features detected).

| Sample name | Sample component | Country of origin | Product | Dilution in Mix | No. of features | Unique features |
| --- | --- | --- | --- | --- | --- | --- |
| PET Mix | PET 1 | Germany | Oven bag | 1:3 | 521 | 846 |
|  | PET 2 | UK | Oven bag | 1:3 | 96 |  |
|  | PET 3 | Norway | Food container | 1:3 | 292 |  |
| PVC Mix | PVC 1 | Germany | Drinking tube | 1:3 | 1374 | 4222 |
|  | PVC 2 | Germany | Drinking tube | 1:3 | 700 |  |
|  | PVC 3 | UK | Cling film | 1:3 | 3804 |  |
| Assorted Mix | PE 1 | Germany | Zip lock freezer bag | 1:8 | 1202 | 5578 |
|  | PE 2 | S. Korea | Freezer bag | 1:8 | 229 |  |
|  | PE 3 | USA | Cling film | 1:8 | 356 |  |
|  | PP 1 | S. Korea | Coffee cup | 1:8 | 642 |  |
|  | PP 2 | USA | Yoghurt cup lid | 1:8 | 990 |  |
|  | PS 1 | USA | Cup<sup>a</sup> | 1:8 | 1534 |  |
|  | PS 2 | Norway | Tray<sup>a</sup> | 1:8 | 526 |  |
|  | PUR 1 | Germany | Hydration bladder | 1:64 | 3050 |  |

Note: <sup>a</sup> = extruded polystyrene, S. Korea = South Korea, UK = United Kingdom of Great Britain and Northern Ireland, USA = United States of America

### 2.2 Sample selection and preparation

All 14 extracts were individually produced via a solvent-based extraction as described in depth elsewhere^27^ and subsequently combined to produce the final Mixes. Briefly, 13.5 g plastic of each sample were extracted in 90 mL methanol (99.8%, Sigma-Aldrich) by sonification for 1 h. Without evaporating to dryness, we removed a 60 mL aliquot from the 90 mL extract, added 600 µL dimethyl sulfoxide (DMSO), and evaporated the samples under a gentle stream of nitrogen to a final volume of 600 µL. Four procedural blanks were included to control for potential contamination. The PVC and PET Mixes were then made by combing equal volumes of the three PVC or PET extracts, respectively. The Assorted Mix was made similarly with the combination of eight extracts, however, PUR 1 was diluted 1:8 in DMSO. We present the concentration of plastic as mg plastic well^−1^, which corresponds to the chemicals extracted from that mass of plastic dissolved and analyzed in 60 µL cell culture media per well.

### 2.3 Nontarget chemical analysis

We performed a nontarget chemical analysis of the plastic extracts as part of a previous study with the methods detailed there (corresponding sample names in Table S1).^27^ In brief, we analyzed each sample on a Waters Acquity UPLC I-Class coupled to a quadrupole time-of-flight mass spectrometer (Synapt G2-S HDMS, Waters) run in positive electron spray ionization mode and performed the data analysis using Progenesis QI (Metascope algorithm, Nonlinear Diagnostics) as previously reported.^27^

Since we used a smaller sample set than before,^27^ a re-analysis of the data allowed us to align all samples included here and created a joint list of chemical features (i.e., ions with a unique m/z and retention time). Only features that were absent in the procedural blanks or had an at least 10-fold higher raw abundances were included.

### 2.4 Cell care and plasmid isolation

HTLA cells, HEK293 cells stably expressing a tTA-dependent luciferase reporter and a β-arrestin-TEV protease fusion gene (kindly provided by Brian Roth) were maintained in DMEM (Sigma Aldrich, D6492), supplemented with 10% FBS (Sigma Aldrich, F9665), 2 μg mL^−1^ puromycin (58-58-2, ≥98%, Invitrogen) and 100 μg mL^−1^ hygromycin (31282-04-9, ≥98%, Invitrogen). Cells were split 1:5 when confluency reached 70% and discarded after passage 10. All GPCR plasmids originated from the Roth lab PRESTO-Tango kit (Addgene, Kit #1000000068) and were amplified in *E. coli* and isolated via mini-preps (Promega, A1330). Plasmid stocks for each GPCR had a 260/280 ratio of 1.7–1.9, a 260/230 ratio of ≥1.8, and concentrations of ≥100 ng/µL.

### 2.5 Selection of GPCRs for primary screen

While the PRESTO-Tango assay is designed for simultaneous screening of 315 non-olfactory GPCRs, 180 of those receptors were not validated with an agonist.^22^ Without validation, we cannot differentiate false negatives from true negatives, and those receptors were, thus, excluded. An additional 22 of the validated receptors were excluded because amplification did not produce sufficient plasmid quantity or quality. The total of 126 GPCRs were included in this study (Table S2).

### 2.6 Transfection efficiency and immunostaining

Prior to the screen, we optimized transfection efficiency based on number of seeded cells, volumes of transfection reagents, and plasmid concentrations using six GPCRs (arginine vasopressin receptor 1A (AVPR1A), arginine vasopressin receptor 1A (AVPR1B), arginine vasopressin receptor 2 (AVPR2), human copper transporter 1 (HCTR1), melatonin receptor 1A (MTNR1A), melatonin receptor 1B (MTNR1B) (Figure S2). Transfected HTLA cells were fixed (4% paraformaldehyde, Thermo Scientific, 28908), quenched (0.1 M ammonium chloride, ≥99.5%, Sigma Aldrich, A9434), permeabilized (0.5% Triton-x, Sigma Aldrich, X100), and blocked, all at room temperature. Cells were incubated first with anti-Flag antibody (1:500, polyclonal mouse anti-Flag, Sigma Aldrich, F1804) then with Alexa Fluor 594-conjugated goat anti-mouse antibody (1:200, Invitrogen, A32742) and nuclear dye (Fisher Scientific, R37605). Cells were washed with PBS (0.5 mM CaCl_2_, pH 7.4), sealed, and stored until imaging. Two images per field (NucBlue and Alexa Fluor) and two fields per well were captured on a Cytation 5 cell imaging multimode reader (BioTek) (details in supporting methods and materials). Images were analyzed using the open source software CellProfiler^28^ (Supporting methods and materials). Transfection efficacy was calculated as a ratio between the number of transfected cells and the total number of cells in the well.

### 2.7 Cell viability

Cell viability was measured using the nuclei count data (NucBlue staining) in non-transfected HTLA cells. We defined cytotoxic effects as a 20% reduction of cell viability as compared to the controls. Briefly, cells were seeded at a density of 10 000 cells well^−1^ in white, optical bottom 384-well plates (Sigma Aldrich, CLS3765) coated with 0.1 mg mL^−1^ poly-L-lysine (Sigma Aldrich, P2636). The following day, cells were starved for 1 h in starving media (DMEM supplemented with 1% dialyzed fetal bovine serum (dFBS, ThermoFisher, A3382001), 1x penicillin/streptomycin) before addition of BPA, DEP, TPP, plastic extracts, and Mixes. The exposure lasted for 23 h before staining with NucBlue and imaging, as described above. To confirm that cell viability was similar between transfected cells, we compared non-transfected cells to those expressing MTNR1B, and AVPR2. No differences were observed at concentrations relevant to the screen (Figure S3) and results were generalized to all receptors.

### 2.8 Presto-Tango assay

Both the primary and secondary screens were performed as previously described^29^ with several notable modifications. On day one, cells were seeded as described in 2.7 and transfected 24 h later (day 2) using Lipofectamine 3000 following the supplier’s instructions (ThermoFisher, L3000008). On day three, the transfection media was replaced with starving media and, after 1 h of starvation, the chemicals and Mixes were diluted in starving media and added to the cells (described in the Supporting information). The exposure lasted for 23 h before cells were lysed and quantification of luminescence was done on Cytation 5 cell imaging multimode reader (BioTek).

For the primary screen, the exposure and transfection layout were such that each plate contained ten receptors exposed to 10 µM of BPA, DEP, or TPP as well as 0.9 mg plastic well^−1^ of the PVC, PET, or Assorted Mix, a negative control (starving media only), and solvent control (0.2% DMSO). Additionally, cells transfected with MTNR1B and exposed to melatonin (≥98%, Sigma Aldrich, M5250) served as a positive control on each plate while non-exposed, non-transfected cells served as background (Figure S4). Each control and treatment were analyzed in four technical replicates (i.e., wells) in a single experiment. For the secondary screen, we constructed seven-point dose-response curves from a 1:2 dilution series with the highest concentration being 30 μM for the single chemicals and 1.8 mg plastic well^−1^ for the Mixes. All secondary screen experiments included four technical replicates and three biological replicates (i.e., independent experiments).

### 2.9 Activity of individual plastic extracts

We further investigated which of the FCAs constituting the active Mixes caused the GPCR agonism. Therefore, we analyzed the three extracts used in the PVC Mix and the eight extracts used in the Assorted Mix in the PRESTO-Tango assay for adenosine receptor 1 (ADORA1) and MTNR1A as described above with some modifications. We used 5’-N-Ethylcarboxamidoadenosine (NECA, Abcam, ab120440) and melatonin as positive controls for ADORA1 and MTNR1A, respectively. A negative control (n = 8) and a five-point solvent control (highest solvent concentration = 0.2%) constructed with a 1:2 dilution series was also included on each plate.

### 2.10 Confirmation using pharmacological knock-down

To further confirm that the observed activation was indeed caused by agonism at the respective GPRC, we co-exposed cells to the active samples and to the ADORA1 antagonist 8-Cyclopentyl-1,3-dipropylxanthine (DCPCX, 99.02%, MedChemExpress, HY-100937)^30^ or the MTNR1A antagonist luzindole (97%, ThermoFisher, J61915-#0).^31^ We used the EC_80_ of NECA (0.05 µM) and of melatonin (0.01 µM) as background agonist for DCPCX and luzindole, respectively. Based on the dose-response relationship of the antagonists, we selected three concentrations of DCPCX (8, 32, and 2000 pM) or luzindole (0.1, 1, and 10 µM). Cells transfected with ADORA1 or MTNR1A were then exposed to 1.8 mg plastic well^−1^ of PUR 1, PVC 1, PVC 2, PVC 3 in combination with three concentrations of the respective antagonist (Supporting methods and materials).

### 2.11 Gene Ontology analysis

Using AmiGO2,^32,33^ we extracted all gene ontology (GO) terms annotated to ADORA1 or MTNR1A. We filtered for terms specific to biological processes in mammals. We prioritized and ordered the GO terms using a structure-based ranking with the R package “GOxploreR”.^34^ Briefly, we removed redundant GO terms that were within the same GO directed acyclic graphs (GO-DAGs)(function “prioritizedGOTerms”) and ordered the remaining terms (function “scoreRankingGO”). The provided score (s_t_) was then used to rank terms based on biological specificity. In this manner, we emphasize biological processes where the effects of ADORA1 and MTNR1A disruption may be better understood.

### 2.12 Data analysis and quality control

All data and statistical analyses were conducted in R (R Core Team, 2022) or GraphPad Prism (v10, GraphPad Software, San Diego, CA). Data visualization and clustering was performed in R with ‘pheatmap’, ‘eulerr’, and ‘dendsort’ packages. The dose-response curves were fit with the ‘drc’ package using a four parameter logistic function (lower limit, upper limit, slope, EC_20_).^35^ For the cytotoxicity of the individual extracts and chemicals, the upper limit was constrained to 100 and the lower limit to 0. For all other curves there was no upper or lower limit constraint.

The solvent and negative controls for the primary screen with all GPCRs were not significantly different (Wilcox-Mann Whitney test, p > 0.05) and were pooled as a measure of constitutive activity (CA). The fold activation was calculated as luminescence of exposed cells (n = 4) divided by CA (n = 8). For the experiments with the individual plastic extracts and the antagonist assay, luminescence was normalized to the CA (0%) and the upper limit of the positive control (100%). Z-scores were calculated to account for the standard deviation within replicates and accurately represent depression of activity (equation S1). A centroid hierarchical cluster analysis of all GPCRs and chemicals and Mixes was preformed using the z-scores from the primary screen in the “pheatmap” package.

As there is no defined threshold in the literature of what is considered a hit,^22,29^ we calculated the limit of detection (LOD) for each receptor as three times the standard deviation (SD) plus the mean of the CA (n = 8). We considered a receptor activation >LOD as a hit that warranted further investigation. For the secondary screen, we calculated the LOD from three biological replicates. For the recovery assay, one-way ANOVA was performed in GraphPad Prism.

Quality criteria were applied to the PRESTO-Tango results based on the data obtained from the negative, solvent, and positive controls on each plate. Stricter quality criteria for the secondary screen and antagonist experiments were employed for quality assurance of data (Supporting methods and materials and Table S3).

## 3 Results and discussion

In this work, we performed the first large-scale screen to determine if plastic chemicals can disrupt GPCR signaling by acting as agonists. We found that FCAs made of PVC and PUR contained potent activators of ADORA1 and MTNR1A and confirmed the specificity of these novel receptor-chemical interactions using known GPCR antagonists. Finally, we identified biological processes involved in the targeted GPCRs and discuss potential implications for human health.

### 3.1 Plastic food packaging contains thousands of chemicals

Prior to the primary screen, we investigated the chemical composition of the plastic extracts using nontarget high-resolution mass spectrometry and investigated the cytotoxicity of the samples to determine non-cytotoxic concentrations to be used in the primary screen.

In total, we detected 10 646 chemical features across all extracts (Table 1). In the individual samples, the number of features ranged from 96 (PET 2, oven bags) to 3804 (PVC 3, cling film). Of the three plastic Mixes, the Assorted Mix contained most features (5578), with 54% originating from PUR 1 (hydration bladder, 3050). PUR 1 was also the most cytotoxic extract (EC_20_ = 0.05 mg plastic well^−1^) and, thus, had to be diluted by an additional factor of 1:8 in the Assorted Mix. The other seven extracts in that Mix contained fewer features and their cytotoxicity was not associated with a specific polymer type or the number of features (Figure S5A). The PVC Mix contained 4222 chemical features, with 90% coming from PVC 3 (cling film), which was also the most cytotoxic PVC extract (EC_20_ = 2.8 mg plastic well^−1^). The PET Mix contained 846 features, with large differences between similar products (PET 1 = 521 vs PET 2 = 96). Despite these differences, the cytotoxicity of all three PET products were similarly low. As expected, the Mixes were more cytotoxic than their individual components. As each individual component was diluted, this is likely due to the larger numbers of chemical features resulting from combining the extracts. Indeed, cytotoxicity increases with the number of chemicals in a sample.^36^ Congruent with previous reports,^26,37,38^ our results demonstrate the presence of a large number of chemicals in plastic FCAs that induce cytotoxicity, particularly in case of PVC and PUR products.

Of the single plastic chemicals included in this study, DEP (EC_20_ = 12.8 µM) and TPP (EC_20_ = 15.0 µM) were most cytotoxic (Figure S5B). BPA (EC_20_ = 34.6 µM) was not cytotoxic up to 30 µM, however at higher concentrations also showed similar suppression of cell viability to that of DEP and TPP (Figure S6D).

### 3.2 Plastic chemicals activate GPCRs

The primary screen was used as a “first pass” to identify receptor-chemical interactions. In total, we tested 756 potential interactions and found eleven hits on GPCRs from the adenosine, melatonin, apelin, lysophospholipid, melanocortin, prostanoid, 5-hydroxytryptamine, and vasopressin families (Figure 1A). These hits activated the respective receptor with a fold change of 1.61–4.22 and z-scores of 3.04–8.88 (Table S4). Unsurprisingly, hits were more prevalent and active across the Mixes than the single chemicals (Figure 1A). The PVC Mix activated four receptors and was most effective on MTNR1A followed by the Assorted Mix on ADORA1.

**Figure 1.**
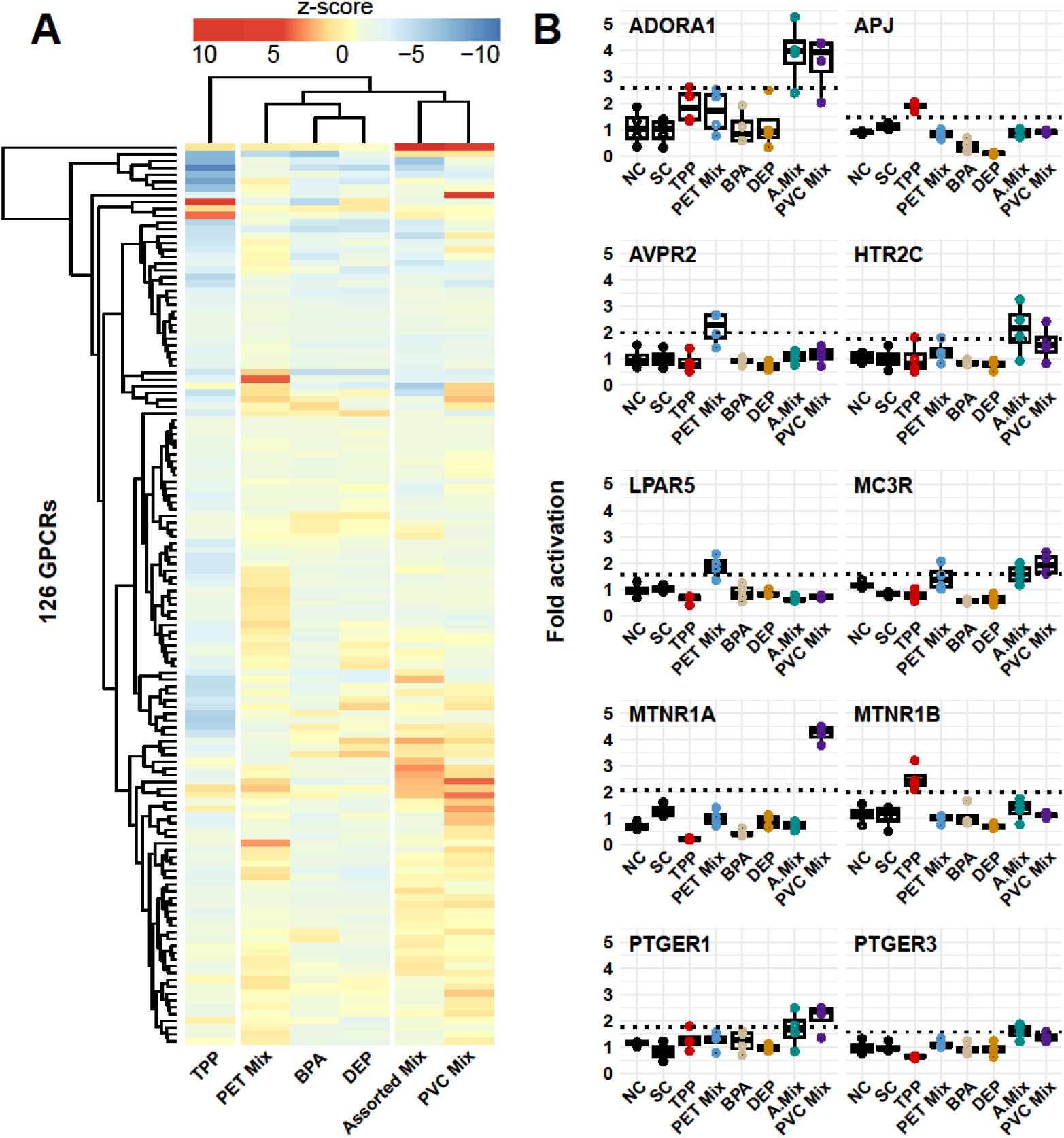
Results of the primary PRESTO-Tango screen of 126 GPCRs exposed to triphenol phosphate (TPP), PET Mix, bisphenol A (BPA), diethyl phthalate (DEP), Assorted Mix, and PVC Mix. (A) Heatmap displaying z-scores where red and blue indicate receptor activation and suppression, respectively. GPCRs, chemicals and Mixes are clustered based on activity. (B) Individual hits from primary screen were determined based on a receptor activation >LOD (dotted black line), the inactive chemicals and Mixes are presented for comparison. NC = negative control, SC = solvent controls, n = 4 (technical replicates).

In contrast, TPP inhibited the activity of 89% of the GPCRs (Figure 1A). Such universal reduction in luminescence across many diverse GPCRs points towards a generic inhibitory effect rather than inverse agonism or antagonism, although experimental validation is needed. One cause could be an inhibition of the luciferase reporter which is a well-known issue in drug screening that increases the probability of false-negative hits.^39^ Interestingly, despite this inhibitory effect, TPP was the only single chemical that activated any GPCR above the LOD (MTNR1B, fold activation = 2.52, z-score = 4.32).

BPA and DEP did not produce a hit at any of the 126 GPCRs suggesting they do not act as agonists of the receptors investigated here. This is somewhat contrary to our expectations as previous research has demonstrated that BPA activated ADORA1, dopamine receptor 1 (DRD1), and serotonin receptor 2C (5HT2C).^40^ GPER activation by BPA has also been well established,^5,6,7,41^ but given that GPER was not be validated with an agonist in the PRESTO-Tango screen,^22^ we did not include it here. More likely, the observed inactivity is due to limitations in the study design. For example, the assay may not be sensitive enough to detect partial agonism. As the Tango assay relies on beta-arrestin recruitment as final step of signal amplification, higher receptor density or prolonged receptor signaling is needed to detect agonism, which may not have been achieved with all GPCR-chemical interactions.^42^ Indeed, some receptor responses by a known agonist were as low as 1.3-fold activation,^22^ although ToxCast21 suggests a signal-to-noise ratio of at least 3-fold for high-throughput screens.^40^ This implies that the rate of false-negative hits might indeed be relatively high when using the PRESTO-Tango system in a toxicological setting.

In sum, the primary screen indicates that chemicals present in a range of plastic FCA may activate certain GPCRs. To confirm the robustness and biological activity of these hits, we performed a secondary screen applying a dose-response design.

### 3.3 ADORA1, MC3R, MTNR1A, and MTNR1B are confirmed targets of plastic chemicals

Of the eleven hits, we confirmed four receptor-chemical interactions in the secondary screen: the PVC Mix activated ADORA1, and MTNR1A in a dose-dependent manner, the Assorted Mix did the same at ADORA1, and TPP is an agonist of MTNR1B (Figure 2). This is, to the best of our knowledge, the first work demonstrating that real-world plastic products contain GPCR agonists activating these receptors. The activity of TPP on MTNR1B receptor has also not been previously reported.

**Figure 2.**
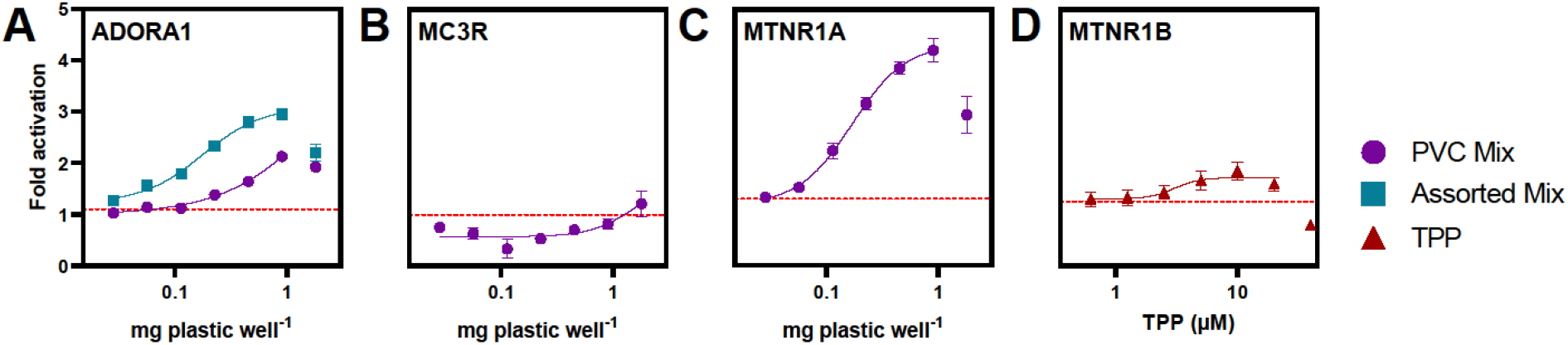
The secondary PRESTO-Tango screen confirms four (A,C,D) and tentatively confirms one (B) out of 11 GPCR-chemical interactions. The highest concentration was in some cases excluded from the dose-response curves due to its sharp decrease in activity. The red horizontal line represents the LOD. Data are shown in mean fold activation ± SEM of three biological replicates with four technical replicates, each.

Chemicals present in the Assorted Mix activated ADORA1 more effectively (3-fold change, EC_20_ = 0.07 mg plastic well^−1^) as compared to the PVC Mix (2.3-fold change, EC_20_ = 0.56 g mg plastic well^−1^, Figure 2A). In contrast to many of the GPCRs investigated here, ADORA1 has been previously screened as a target of exogenous chemicals.^40^ Using three radioligand binding assays, ToxCast identified 115 chemicals as active ADORA1 ligands, including certain phenols and phthalates found in plastics.^40^ In fact, 45 of these are plastic-related substances, some of which have known endocrine activity (Table S5).^18^ However, we did not detect these plastic-related chemicals in our samples,^27^ suggesting that the observed ADORA1 agonist(s) are previously unidentified.

The compounds in the PVC Mix also induced MC3R activity >LOD at 1.8 mg plastic well^−1^ (Figure 2B). While this indicates that MCR3 agonists are present, we were unable to test higher concentrations due to cytotoxicity. Such limitations are common when working with complex mixtures as they can incur nonspecific cytotoxicity before any specific effect can be observed.^36^ Due to the weak response, we were only able to tentatively confirm the hit and did not conduct further experiment. Although we cannot draw conclusions concerning the dose-dependence of the effect of the chemicals present in PVC, we confirm that they activate MC3R at high sample concentrations.

The chemicals in the PVC Mix induced a 4.2-fold activation of MTNR1A with an EC_20_ of 0.08 mg plastic well^−1^ (Figure 2C). Within the same GPCR family, TPP caused a 2-fold activation at MTNR1B (EC_20_ = 2.19 µM), though, the activity decreased at higher concentrations (Figure 2D). Curiously, PVC Mix and TPP did not act similarly on the two melatonin receptors, despite they share the same endogenous ligands and have conserved orthosteric binding sites.^43^ Each receptor selectively binds unique ligands^44^ and differentially recruits beta-arrestin for the same compound,^45^ suggesting the chemicals in this study may act as receptor-selective agonists. While plastic chemicals have not previously been described to target MTNR1A or MTNR1B, several carbamate insecticides with high structural similarities to melatonin agonize or antagonize both receptors.^10,11^

We did not confirm the following hits as they did not produce a dose-response relationship in the secondary screen: PVC Mix at prostaglandin E receptor 1 (PTGER1), Assorted Mix at prostaglandin E receptor 3 (PTGER3), PET Mix at AVPR2, PET Mix at lysophosphatidic acid receptor 5 (LPAR5), and TPP at Apelin receptor (APJ, Figure S7 A-F). The decreasing relationship for certain interactions (Assorted Mix at PTGER3) suggests nonspecific effects are partially or fully masking activation of the receptor, as discussed above in the case of MC3R. In other cases, activation may be caused by a weaker secondary response. For example, the activity of the Assorted Mix on PTGER1 (Figure S7C) may be an indication of a general cellular stress response producing endogenous prostaglandins which then act on PTGER1 in a para-or autocrine manner.^46^

Although only four of the hits from the primary screen could be validated, a subset, particularly ADORA1 and MTNR1A, exhibited robust confirmation. While high throughput assays are susceptible to false positives,^47^ parallel screening facilitates identification of frequent hitters, leaky luciferase expression, and independent activation of the reporter.^22,29^ Therefore, the unconfirmed hits in the present study warrant further investigation to ascertain whether they are true false positives or a sign of more complex receptor-chemicals interactions.

### 3.4 PVC and PUR products contain potent and effective GPCR agonists

Given that the Assorted and PVC Mixes produced the most robust hits in the secondary screen, we investigated the activity of the individual extracts constituting those Mixes. Accordingly, we tested the three extracts in the PVC Mix on ADORA1 and MTNR1A and the eight extracts in the Assorted Mix on ADORA1. In this manner, we identified specific products and polymer types that contained the respective GPCR agonists.

Of the six polymer types included in our study, only products made of PVC and PUR contained GPCR agonists that induced dose-dependent activity (Figure 3). For ADORA1, PUR 1 (a drinking tube) was, by far, the most potent and effective extract inducing a 104% activation at the highest non-cytotoxic concentration (EC_20_ = 0.005 mg plastic well^−1^, Figure 3A–D). Since PUR 1 was the only active extract in the Assorted Mix, it is the source of the ADORA1 agonist(s). However, the lower maximum activity of the Assorted Mix compared to PUR 1 (Figure 3A) also points towards the presence of chemicals in the other extracts that suppress this agonistic effect. Accordingly, receptor activity can be masked by GPCR antagonists or other inhibitors present in plastics. This demonstrates mixture toxicity plays a critical role when testing complex mixtures; an aspect which becomes even more complex considering GPCRs mediate signals both orthosterically and allosterically, often in tandem.^48^

**Figure 3.**
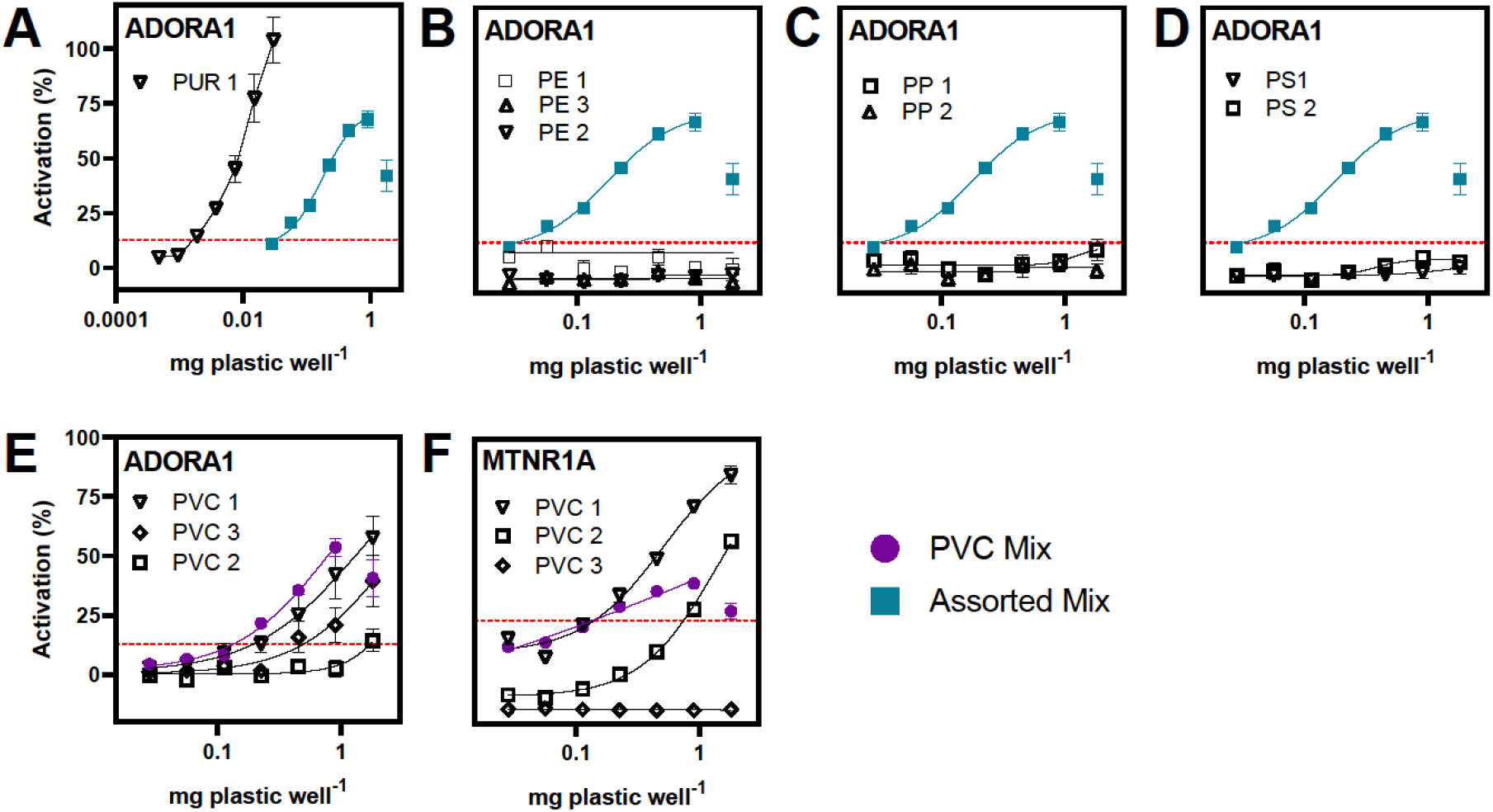
Activity of individual plastic extracts used in the Assorted Mix (blue) on ADORA1 (A–D) and in the PVC Mix (purple) on MTNR1A (E, F). The concentrations of the Assorted and PVC Mix correspond to 1/8 and 1/3 of the concentrations of individual extracts, respectively. Data are shown as mean ± SEM of three biological replicates with four technical replicates, each, and were normalized to the maximal activity of the reference compound and the CA of the receptor. The red dotted line denotes the limit of detection.

Chemicals present in all three PVC products activated ADORA1, albeit with differing efficacies and potencies (Figure 3E). PVC 1 was most active (58%, EC_20_ = 0.35 mg plastic well^−1^), followed by PVC 3 (40%, EC_20_ = 0.79 mg plastic well^−1^), and PVC 2 (14%, EC_20_ = 2.18 mg plastic well^−1^). This shows that ADORA1 agonists are abundant in PVC products. However, given the complexity of the Mixes, we cannot conclude whether the same active chemicals are present in all samples in different concentrations, or whether different chemicals caused the observed activity.

At the MTNR1A, PVC 1 was also most active (85%, EC_20_ = 0.18 mg plastic well^−1^) followed by PVC 2 (57%, EC_20_ = 0.52 mg plastic well^−1^). Interestingly, PVC 3 reduced the activity to approximately 20% below CA at all concentrations (Figure 3F), suggesting the presences of antagonists or other inhibitors in the extract. This is further supported by the comparably lower activation caused by the PVC Mix than by its individual components.

The screening of individual plastic FCAs enables the identification of polymer types containing GPCR agonists and may provide a solution to minimize human exposure to such chemicals, regardless of their identity. For example, it is well established that PUR and PVC are the most problematic polymers due to the use of hazardous chemicals^49^ that induce of a wide array of toxic effects, including endocrine disruption.^26,38,50^ Nonetheless, both materials are still used in some FCA, such as water pipes and, more commonly, in consumer products, including children’s toys.^26^ This study reinforces the existing evidence that PUR and PVC plastics are chemically problematic and should be substituted with safer alternatives.

### 3.5 Confirmation of receptor specificity

To further confirm the receptor specificity of the activity induced by plastic chemicals, we co-exposed HTLA cells with the active PVC and PUR extracts and with known antagonists for ADORA1 and MTNR1A. DCPCX significantly reduced the ADORA1 activity induced by the reference compound as well as PUR 1 and all PVC extracts in a dose-dependent manner in line with their potency (Figure 4A). Similarly, the MTNR1A antagonist luzindole suppressed the activity of PVC 1 and 2 in a dose-dependent manner (Figure 4B). The highest concentration completely knocked down the effect of both extracts and, again, the more potent PVC 1 extract required higher concentrations of luzindole to reduce its activity. These experiments demonstrate that the well-known ADORA1 and MTNR1A antagonists are competing for the same receptor binding sites as the compounds present in PVC and PUR products. We, thus, further confirm that these products contain melatonin or adenosine receptor agonists.

**Figure 4.**
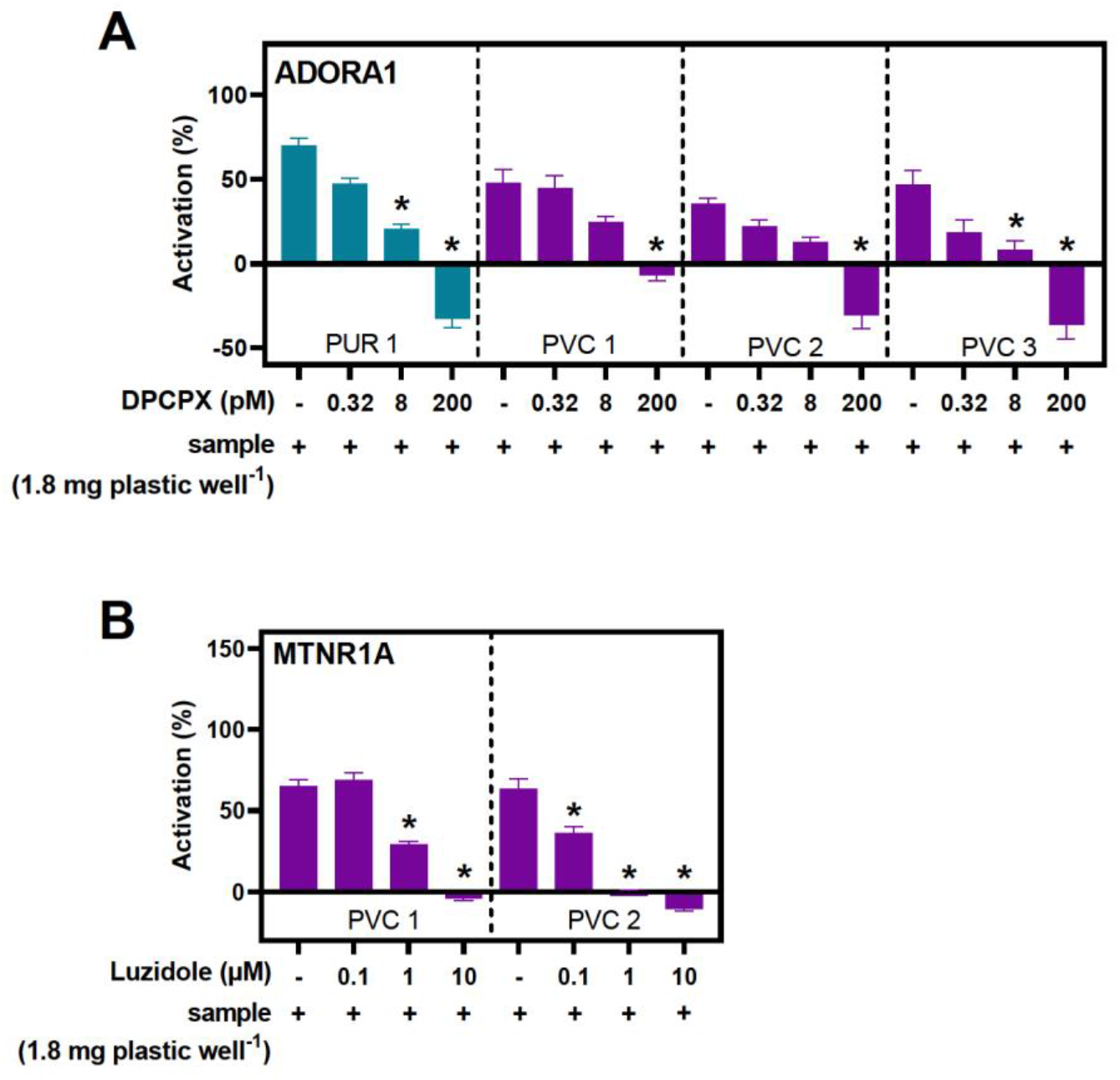
Chemicals in PVC and PUR are competing for binding sites at the (A) ADORA1 and (B) MTNR1A with known receptors antagonists. Co-exposure with DCPCX or luzindole significantly reduces activity of ADORA1 and MTNR1A, respectively. Data are shown as mean ± SEM of three biological replicates with four technical replicates, each, and were normalized to maximal activity of the reference compound and the CA of the receptor. Asterisk indicate significantly different activity from the control (p < 0.05, one-way ANOVA between the control (-)).

### 3.5 Biological implications

Given that plastic chemicals are binding and activating GPCRs, we strove to understand the implications for GPCR signaling and, further down the line, the potential impacts on human health. To do so, we used biological process GO terms annotated to ADORA1 and MTNR1A. The relevance of each annotation was determined by its biological specificity, as quantified by their hierarchical level in relation to the maximal attainable level within the GO-DAG. Given that higher level GO-terms relate more specific biological information,^34,51^ this allows for a more precise characterization of a gene’s function within that process, and therefore, more accurate interpretations of potential biological consequences of GPRC activation.

ADORA1 had 45 annotations after elimination of GO-terms that shared GO-DAGs (Figure S10, Table S6). These terms describe a broad spectrum of biological processes, underscored by the receptors extensive distribution and distinct functionality within various tissues.^52^ Conversely, MTNR1A had five unique annotations. The specific GO term “negative regulation of circadian sleep/wake cycle, non-REM sleep” (s_t_, 0.53) annotated to ADORA1 converges into the same GO-DAG of “circadian rhythm” (s_t_, 0.02), annotated to MTNR1A. Indeed, both ADORA1^53^ and MTNR1A^54,55^ regulate and maintain circadian systems that align cellular, physiological, and behavior processes into a 24-hour cycle. Disruption of ADORA1 signaling is linked to alterations in sleep,^56,57^ an effect most people have experienced through the consumption of caffeine, an ADORA1 antagonist.^58^ Likewise, melatonin or its synthetic mimics, are commonly prescribed to promote sleep via MTNR1A.^54^ Interestingly, exogenous melatonin can also promote sleep via adenosine receptors by inducing endogenous adenosine production.^59^ This exemplifies the importance of considering mixture toxicity as multiple chemicals, with distinct biological targets, may regulate the same process to amplify disruption within an organism. While our work concerns only molecular initiating events, alterations of circadian homeostasis are clearly linked to serious human health impacts, including cancer,^60^ infertility,^61,62^ and impairment of immune function.^63^

The most specific GO term annotated to MTNR1A was “regulation of insulin secretion” (s_t_, 0.53). In addition to circadian regulation, MTNR1A receptor signaling has been associated with insulin sensitivity and the accumulation of fat, contributing to obesity and diabetes.^64,65^ Along the same lines, ADORA1 is largely expressed in white adipose tissue and regulates triglyceride homeostasis, fatty acid homeostasis, and lipid catabolic processes (Figure S10, Table S6). We have previously shown that chemicals in PVC and PUR products increase adipocyte size and triglyceride content via an unknown PPARγ-independent mechanism.^37^ Given that the stimulation of ADORA1 in white adipose tissue increases adipogenesis,^66^ this could potentially represent a new mechanism that mediates the metabolism-disrupting effects of chemicals in plastics.

These results can be used to generate hypotheses on the downstream effects of plastic chemical disruption of ADORA1 or MTNR1A. However, such inferences are substantially complicated by the complexity of GPCR signaling. For instance, stimulation of ADORA1 in the brain reduces body weight and lipolysis^66^ but has the opposite effect in adipocytes. Although acting on the same receptor, an organ-specific distribution of both the receptor and chemical could elicit contrasting adverse outcomes. As another example, prolonged exposure to MTNR1A agonists can reduce receptor density resulting in antagonistic effects.^67^ In addition, bias signaling, homo-or heterodimerization, and allosteric binding^68^ must be given consideration when extrapolating from in vitro to in vivo systems. Nevertheless, the potential link between metabolic and circadian disruption mediated through ADORA1 and MTNR1A warrants follow-up research.

### 3.6 Limitations and future directions

In this work, we show that plastic FCAs contain potent GPCR agonists using multiple layers of evidence (replication, dose-dependency, pharmacological knock-down). This is significant because, historically, research has largely focused on chemicals acting via nuclear receptors^69^ and overlooked GPCRs as targets. Our work addresses this blind spot and highlights the need to expand our focus to include a broader range of receptors.

While primarily designed for drug discovery, the PRESTO-Tango assay is a powerful tool for toxicological research However, given that drug screens are designed to search for very potent agonists, it likely lacks the sensitivity to detect compounds with weak GPCR activity. While this work focused exclusively on agonist activity at GPCRs, we found evidence for the presence of antagonists within the plastic samples that may mask agonist activity thereby increasing the rate of false negatives. Certainly, it is possible to optimize the assay for specific receptors of interest, but our main aim was to identify the most robust GPCR-chemical interactions across many receptors in a high-throughput, simultaneous fashion.

In this manner, we were able to show that chemicals in plastic food packaging act as agonists of ADORA1 and MTNR1A. Three lines of follow-up research arise from this: Firstly, we have investigated all extractable chemicals from plastic FCAs using methanol. To investigate the potential of human exposure more closely, migration studies using more realistic food simulants, such as water or ethanol, can help to assess the leachability of GPCR agonists from plastics. Secondly, effect-directed analyses can be applied to identify the compounds causing the ADORA1 and MTNR1A activity. While this is not without challenges, such identification would facilitate replacing the active chemicals in PUR and PVC. Thirdly, an in vivo assessment of the effects on circadian and metabolic processed caused by ADORA1 and MTNR1A activation is required to understand how our in vitro findings translate to complex biological systems.

While in an early stage of discovery, our work adds GPCRs to the lists of molecular targets that can be disrupted by plastic chemicals. Given that melatonin is an integral part of the hormone system, MNTR1A disruption illustrates that EDCs can act via cell surface receptors as well. However, GPCRs not directly connected to the endocrine system should not receive less attention as exemplified by the ADORA1 agonists present in plastic. Here, insights from pharmacological research can be used to prioritize GPCRs that are important drug targets for further toxicological research.

In a broader context, this study contributes to the growing evidence that plastic products contain compounds, or mixtures of compounds, that elicit a diverse range of toxic effects. A fundamental reconsideration and redesign of the way we make, and use plastics is imperative if plastics are to be considered safe. By adopting strategies that reduce the number and hazard of chemicals used in plastics, we can minimize exposure and reduce their contribution to the burden of disease.

### Potential competing interests

M.W. is an unremunerated member of the Scientific Advisory Board of the Food Packaging Forum Foundation and received travel support for attending annual board meetings.

## Acknowledgements

We thank Felicity Ashcroft for her expertise on optimization of cell culture assays. We also thank Brian Roth for the donation of HTLA cells. Finally, thanks to Dru Jagger, Jaeho Lee, and Mara McPartland with their help on obtaining our plastic samples. This work was mainly supported by internal funding by the Norwegian University of Science and Technology and in parts by the European Union’s Horizon 2020 research and innovation programme under the Marie Skłodowska-Curie grant agreement No 860720.

## Author contributions

M.M., J.V., and M.W. conceived the study, M.M. wrote the manuscript, M.W., and J.V., reviewed and edited the manuscript, M.M., S.S., and I.G. preformed the plastic sampling and preparation, M.M. and I.G. performed the in vitro experiments, Z.B. and M.W. performed the chemical analysis, M.M preformed the data analysis.

**Supporting information for this study is available on bioRxiv**

