## Supporting information, methods and materials, figures 1 - 10 for "Beyond the nucleus: Plastic chemicals activate G protein-coupled receptors"

[2] present address: Innovative Environmental Services (IES) Ltd, Benkenstrasse 260, 4108 Witterswill, Switzerland

Number of pages: 19

Number of figures: 10

### Contents

The supporting information includes text, tables, and figures detailing:

Supporting methods and materials

Transfection efficiency and immunostaining

Cell profiler analysis

Cell viability

PRESTO-Tango assay

Activity of individual plastic extracts

Data analysis

Supporting Figures

Figure S1. Schematic of items and mixtures that constitute each Mix included in the study

Figure S2. Receptor expression for a selection of GPCRs

Figure S3. Cell viability of non-transfected cells and transfected cells.

Figure S4. Cytotoxicity of plastic extracts, Mixes, and bisphenol A (BPA), diethyl phthalate (DEP), and triphenol phosphate (TPP)

Figure S5. Cytotoxicity of non-transfected HTLA cells

Figure S6. Plate layout for the PRESTO-Tango primary screen

Figure S7. The secondary PRESTO-Tango screen hits that could not be confirmed in a dose dependent manner

Figure S8. Dose-response relationship for (A) ADORA 1 exposed to reference compound NECA, (B) MTNR1A and (C) MTNR1B exposed to reference compound melatonin

Figure S9. Dose-response relationship for (A) ADORA1 and (B) MTNR1A exposed to reference antagonists DCPCX and luzindole

Figure S10. Biological processes regulated by ADORA1 and MTNR1A

### Supporting Methods and Materials

**Transfection Efficiency and immunostaining**

Transfected HTLA cells were fixed (4% paraformaldehyde, Thermo Scientific, 28908) for 20 min, quenched with 0.1 M ammonium chloride (≥99.5%, Sigma Aldrich, A9434) for 10 min, permeabilized with 0.5% Triton-x (Sigma Aldrich, X100) for 30 min, and blocked with 5% powdered milk for 1 h, all at room temperature. Cells were then incubated with anti-Flag antibody (1:500, polyclonal mouse anti-Flag, Sigma Aldrich, F1804) overnight at 4 °C. The following day, we incubated the cells with Alexa Fluor 594-conjugated goat anti-mouse antibody (1:200, Invitrogen, A32742) and nuclear dye (Fisher Scientific, R37605) for 1 h in the dark at room temperature. Cells were thoroughly washed with PBS (0.5 mM $CaCl_2$, pH 7.4), sealed, and stored in the dark at 4 °C until imaging.

Fluorescence per well was assessed using a Cytation 5 Cell Imaging Multimode reader (BioTek) with excitation at 485 nm and emission at 572 nm for Alexa Fluor, and excitation at 360 nm and emission at 460 nm for NucBlue. Imaging took place on the same instrument utilizing a 10x Plan Fluorite objective (WD10, NA 0.3). NucBlue fluorescence was selected as the image plane through image-based autofocusing, and two images per field were captured

(NucBlue and NileRed). Detection of NucBlue staining was accomplished using a 365 LED with DAPI filter cube (Ex 377/50, Em 447/60), while Alexa Fluor was detected with a 523 LED using an RFP filter cube (Ex 531/40, Em 593/40).

**Cell Profiler analysis**

For transfection efficiency, Alexa Fluor and NucBlue staining were imaged at 10x magnification and analyzed using the following protocol to quantify the number of cells in each well (NucBlue) and number of transfected cells in each well (Alexa Fluor).

Alexa Fluor transfection data:

1. Identification of nuclei and transfected cells

Primary objects (nuclei) were identified using a Global threshold strategy and Otus thresholding method from images stained with NucBlue. The primary objects were used as the seed objects to identify the secondary objects (transfected cells). Transfected cells were identified from the Alexa Fluor images using a propagation method with the same thresholding strategy and method. The intensity of the primary (nuclei) and secondary (transfected cells) objects were subsequently measured.

*2. Data processing*

First, filtering was applied to identify any nuclei (which were then outlined and overlayed on the NucBlue images). The identified nuclei were overlayed on the Alexa Fluor image to identify the transfected cells. A calculation between overall cells in the image (NucBlue) and number of transfected cells in the corresponding image (Alexa Fluor) was used to calculate transfection efficiency.

An additional pipeline was used to quantify cytotoxicity by imaging NucBlue staining images at 4x magnification.

Nuclei count data:

*1. Identification of dead cells in the foreground*

Dead nuclei fluorescing more brightly and in the foreground of the image were identified using an Adaptive Otsu thresholding method with two classes.

*2. Enhancing nuclei*

The neurite features of original image were then enhanced using the line structures method. This enabled less brightly fluorescing cells to be more clearly identified.

*3. Identification of all cells*

Using the enhanced image, all nuclei were identified using an Adaptive strategy Otsu threshold method now with three classes. We then overlayed counted the dead cells identified in step 1 and subtracted them from all the cells counted in step 3 to obtain final nuclear counts.

#### Cell viability

NucBlue staining was used to obtain nuclei count data in non-transfected HTLA cells. Cell viability was then defined as a 20% of cells (nuclei) as compared to controls. On the day of exposure (day 2) media was removed and replaced with 30 µL starving media (DMEM supplemented with 1% dialyzed fetal bovine serum (dFBS, ThermoFisher, A3382001), 1x penicillin/streptomycin) for 1 h. During this time exposure media containing BPA, DEP, TPP, plastic extracts, and plastic Mixes were prepared. For BPA, DEP, and TPP stock concentrations of 20 mM were prepared and then diluted 1:240 into starving media to make the highest concentration of 480 µM. Serial 1:2 dilutions were then performed to create 7 concentrations for the dose–response curves. Finally, the exposure media was further diluted 1:2 into the well (addition of 30 µL into the existing 30 µL in the well) for the concentration range from 240–3.75 µM.

For the individual plastic extracts and plastic Mixes the stock concentration contained 900 mg plastic well$^{-1}$, or 15 000 g plastic/L. From this stock we diluted 1:260 and again make 1:2 serial dilutions to create 8 concentrations. There was another 1:2 dilution step into each well (addition of 30 µL into the existing 30 µL in the well) for the concentration range from 22.5–0.17 mg plastic well$^{-1}$. The exposure lasted for 23 h before two washing steps with PBS, staining with NucBlue for 30 min, and imaging, as described above.

**Presto-Tango assay**

The primary and secondary screens follow the detailed protocol provided by Zeghal et al. (2020)[1]. On day one, cells were seeded as described in 2.7 and transfected 24 h later (day 2). Prior to transfection, the media was replaced with 30 µL growth media. Lipofectamine 3000 was used to ensure the best transfection across receptors, though this deviates from the transfection protocol described by Zeghal et al. (2020)[1]. Transfection media (10 µL) was mixed in a 96-well plate, where each well contained 200 ng of a unique GPCR plasmid. Well A1 an A12 always contained MTNR1A plasmid and TE buffer, respectively. Transfection media was then added to each well and gently tapped to ensure all transfection media was in the bottom well. On day three, the transfection media (total of 40 µL) was replaced with starving media (30 µL) and an additional 30 µL exposure media was added to the cells. For BPA, DEP, and TPP stock concentrations of 20 mM were diluted 1:1000 into starving media to make the concentration of 20 µM. The exposure media was added to the cells in another 1:2 dilution step (addition of 30 µL into the existing 30 µL in the well) for the final concentration 10 µM. For Assorted, PVC and PET Mixes stock concentrations of 900 mg plastic well$^{-1}$ (15 000 g plastic/L) were diluted 1:500 to make the concentration of 1.8 mg plastic well$^{-1}$. Again, another 1:2 dilution step (addition of 30 µL into the existing 30 µL in the well) resulted in the final concentration of 0.9 mg plastic well$^{-1}$. For the background and solvent control 30 µL starving media with DMSO (0.2%) was added while the negative control contained only starving media.

For the secondary screen, we constructed seven-point concentration-response curves from a 1:2 dilution series with the highest concentration being 30 µM for the single chemicals and 1.8 mg plastic well$^{-1}$ for the mixes. In more detail, BPA, DEP, and TPP stock concentrations of 20 mM were diluted 1:500 into starving media to make the highest concentration of 60 µM. Serial 1:2 dilutions were then performed to create 7 concentrations for the dose-response curves. Finally, the exposure media was further diluted 1:2 into the well (addition of 30 µL into the existing 30 µL in the well) for the concentration range from 30–0.47 µM. For the plastic Mixes the stock concentration containing 900 mg plastic well$^{-}$ were diluted 1:250 and again make 1:2 serial dilutions to create 8 concentrations. There was another 1:2 dilution step into each well (addition of 30 µL into the existing 30 µL in the well) for the concentration range from 1.8 – 0.03 mg plastic well$^{-1}$.

For both the primary and secondary screen MTNR1A activated by melatonin was used as a positive control with a dose-response concentration range of 10–0.00001 µM. Similar to the plate layout for the primary screen, wells K1, L1, K2, L2 served as a solvent control (DMSO, 0.2%) with which activation was calculated against.

After the 23h of exposure the media was removed, a white sticker was placed on the bottom of the transparent plates, and cells were lysed in 20 µL cell lysis buffer (25 mM pH 7.8 TRIS, 2 mM DDT, 2 mM 127 CDTA, 10% glycerol, and 1% Tritron-X100) and linearly shook for 3 minutes. Luminescence was measured (Cytation 5) for one second after injection of 30 µL illuminate mix (20 mM Tricine, 1.07 mM $C_4H_2Mg_5O_{14}$, 2.67 mM $MgSO_4$ x $7H_20$, 0.1 mM EDTA, 1.5 mM DDT, 539 µM D-Luciferine, 5.49 mM ATP) followed by quenching of the reaction with 30 µL 0.1 M NaOH.

**Activity of individual plastic extracts**

To determine which extracts were active in the Mixes we tested the three PVC Mix extracts (PVC 1, PVC 2 PVC 3) against MTNR1A and ADORA1 and the eight Assorted extracts (PE 1, PE 2, PE 3, PS 1, PS 2, PP 1, PP 2, PUR 1) against ADORA1. The total concentration of plastic in the Mixes was calculated as an addition of the three or eight extracts that comprised it. Therefore, where testing the individual extracts, all chemicals have a 3x or 8x higher concentration of the *chemicals* they comprise. However, the overall concentration of *plastic* in each well is the same, so although the chemicals are up concentrated, the mg plastic well$^{-1}$ are comparable between the Mixes and their individual components.

Cells were seeded and transfected as described above. On day 3, exposure was done with the extracts, however now we also included a positive control and reference compound specific for MTNR1A (≥98%, Sigma/Merk, M5250) and ADORA1 5'-N-Ethylcarboxamidoadenosine (NECA, Abcam, ab120440) to be able to quantify activation against a canonical agonist. Melatonin (stock 20 mM) was diluted 1:1000 (20 µM) and serial dilution (1:100) were done to create the concentration range to 10–1 x $10^{-11}$ µM. NECA (stock 10 mM) was diluted 1:100 (100 µM) and serial dilutions (1:10) were done to create the concentration range 50–5 x $10^{-5}$ µM. For the extracts, excluding PUR 1, stock concentration containing 900 mg plastic well$^{-1}$ were diluted 1:250 and 1:2 serial dilutions were done to create seven concentrations. There was another 1:2 dilution step into each well (addition of 30 µL into the existing 30 µL in the well) for the concentration range from 1.8 - 0.03 mg plastic well$^{-1}$. For PUR 1, due to cytotoxicity, the stock solution was diluted 1:16,000, and following the same serial dilution resulted in a concentration range from 0.03 - 0.47–0.00042 mg plastic well$^{-1}$.

For the pharmacological knock-down assay, on each plate there was a positive control with the reference compound (as described above), a seven-point dose-response curve with only the antagonist (DCPCX for ADORA1 and luzindole for MTNR1A), and a seven-point dose-response curve for DCPCX and luzindole combined with background agonist NECA and melatonin (0.01 μM), respectively. The concentration for NECA and melatonin was determined by the $EC_{80}$ of the positive control. For DCPCX without agonist, a 1:5,000 dilution (20 mM stock solution) was made and then 1:5 serial dilutions were done to create seven concentrations from 200–0.01 pM. For DCPCX with agonist, the same was done with the DCPCX however, the dilution was done into starving media previously spiked with NECA (0.05 μM). For luzindole, without agonist, 1:100 dilution (20 mM stock solution) was made and again a 1:5 serial dilution was carried out ending with the concentration range from 100–0.0064 μM. When combing luzindole with melatonin the media was spiked with melatonin (0.01 μM) and the luzindole exposures were made as described above.

From these controls, three concentrations of DCPCX (8, 32, and 2000 pM) and luzindole (0.1, 1, and 10 μM) were selected and mixed with media previously spiked with 1.8 mg plastic well$^{-1}$ of PUR 1, PVC 1, PVC 2, or PVC 3. On day 4, removal of media, cell lysis, and luminescence readings were all carried out as described above.

**Data analysis**

The following quality criteria were applied to the PRESTO-Tango results based on the data obtained for negative, solvent, and positive controls on each plate: The MTNR1B positive control had to produce sigmoidal dose-response curves with a maximum fold induction of ≥40, the $EC_{20}$ had to fall within a range of 0.5–5 mM, the Z' value (equation S1) had to be >0.25 (primary screen) or 0.5 (secondary screen), and coefficient of variance of the $logEC_{20}$ had to be <3% (primary) or 1.5% (secondary screen). The stricter quality criteria for the secondary screen were also applied to the antagonist experiments (Table S3).

**Equation S1.** The Z-score, different from Z', is a statistical measure of how far a value is from the mean, in terms of standard deviation. We used z-score to represent activation in the primary screen to consider the standard deviation of the technical replicates and the depression of activity observed in many interactions. Z-score is calculated using the observed value (*x*), the mean (μ) of the replicates, and the standard deviation (σ) from calculating that mean.

$$z\text{-}score = \frac{x - \mu}{\sigma}$$

**Equation S2.** Z' was used to measure the quality of the positive control on each plate in the primary and secondary screen. The calculation uses the standard deviation of the upper limit of the positive control ($\sigma_{top}$), the standard deviation of the bottom limit of the positive control ($\sigma_{bottom}$), the upper limit of the positive control ($\mu_{top}$), and bottom limit of the positive control ($\mu_{bottom}$). A Z' between 0.5 and 1 is considered excellent, although <0.5 is also acceptable in biological assays[2].

$$Z' = 1 - \frac{3\,(\sigma_{top} + \sigma_{bottom})}{|\mu_{top} - \mu_{bottom}|}$$

### Supporting Results

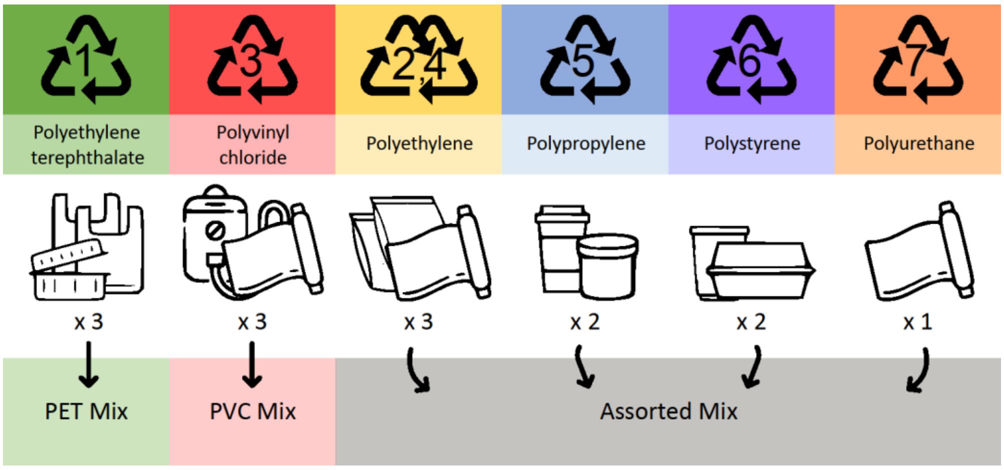


**Figure S1. Schematic of items and mixtures that constitute each Mix included in this study**.

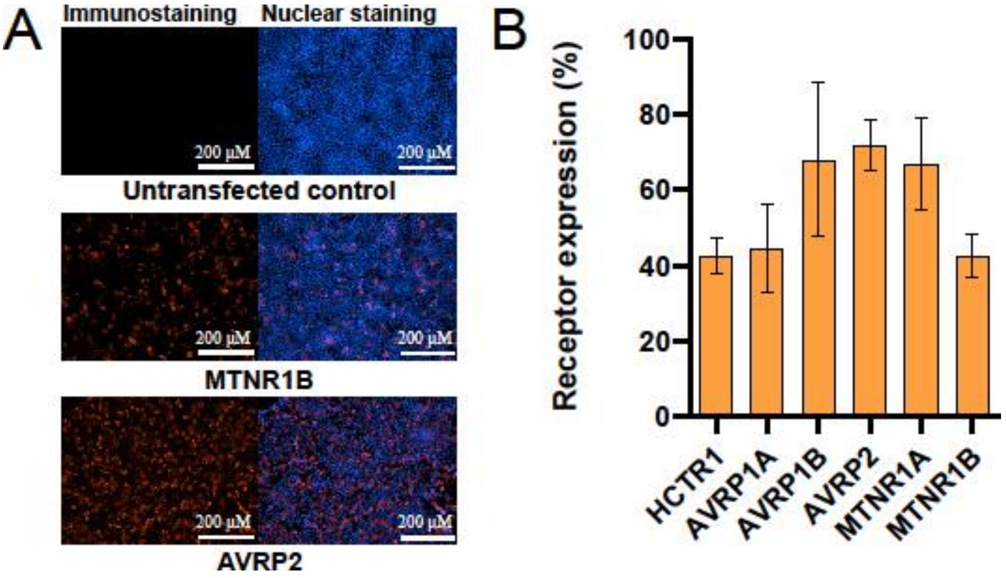


**Figure S2. Receptor expression for a selection of GPCRs included in the primary screen and used to optimize the transfection procedure.** (A) Surface expression of AVRP2 and MTNR1B as shown by immunofluorescence with an anti-flag antibody with an overlay of nuclear staining. (B) Bar chart comparing receptor expression between MTNR1B, HCTR1, AVPR1A, MTNR1A, AVPR1B, and AVPR2 as percent of cells expressing the respective GPCR.

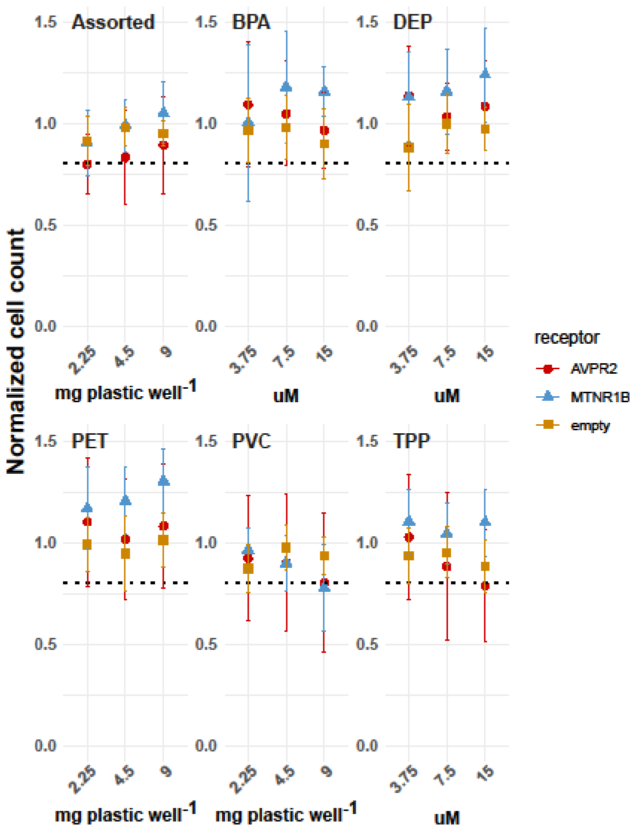


**Figure S3. Cell viability of non-transfected cells and transfected cells exposed to plastic mixes or single chemicals**. MTNR1B transfected cells (blue), AVPR2 (red) and non-transfected cells (empty indicating empty vector, orange) exposed to Assorted Mix, BPA, DEP, PET Mix, PVC Mix, and TPP. Concentration (x-axis) is given in mg plastic well$^{-1}$ for Assorted Mix, PET Mix, and PVC Mix and in µM for BPA, DEP and TPP. The black horizontal line represents the cytotoxicity cut-off of 0.8. Data is normalized to the cell count of controls and shown in mean cell count ± SEM of four technical replicates and three biological replicates.

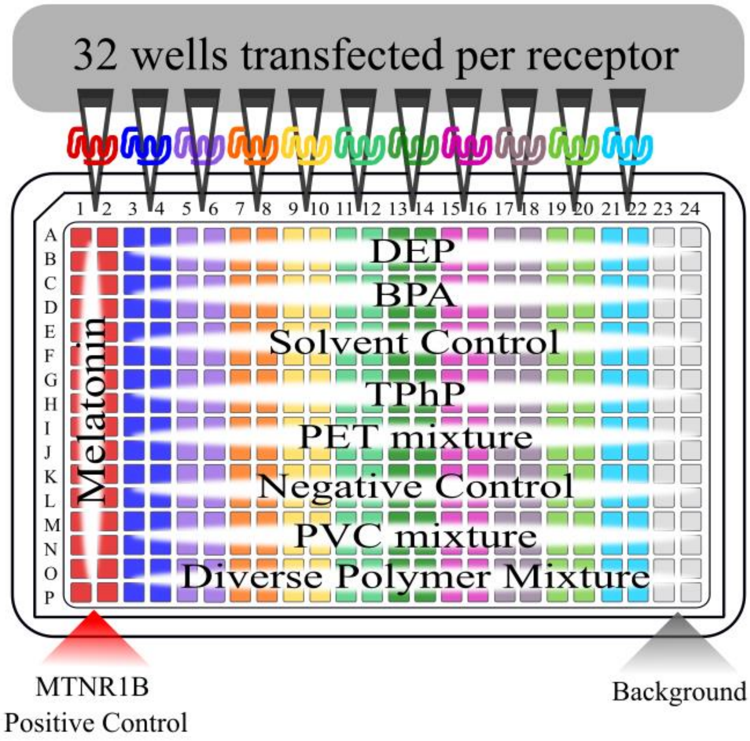


**Figure S4. Plate layout for the PRESTO-Tango primary screen of 126 GPCRs.** The positive control is set up on the right of the plate and background on the left. The different colors indicate different receptors of the 126 that were transfected, and the exposures run horizontally across the plate.

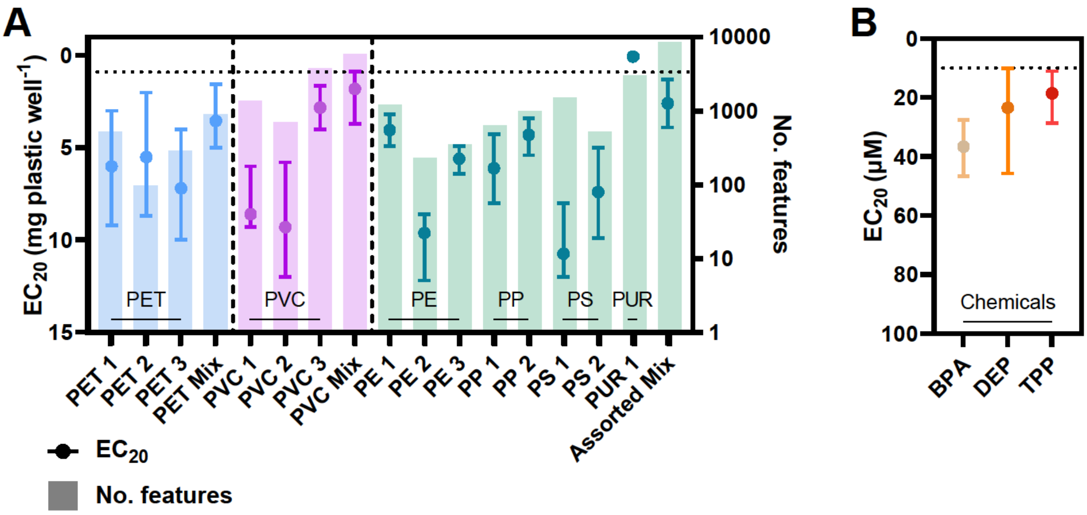

**Figure S5. Cytotoxicity of (A) plastic extracts and the corresponding mixes (grey area) as well as (B) bisphenol A (BPA), diethyl phthalate (DEP), and triphenol phosphate (TPP) in non-transfected HTLA cells.** (A) also contains the number of chemical features detected in each sample. The dotted black line represents the concentration used in the primary screen. Note that PUR 1 was analyzed in an 8-fold lower concentration. The data is presented as mean $EC_{20}$ ± 95% CI for nuclei count data from three biological replicates with four technical replicates, each.

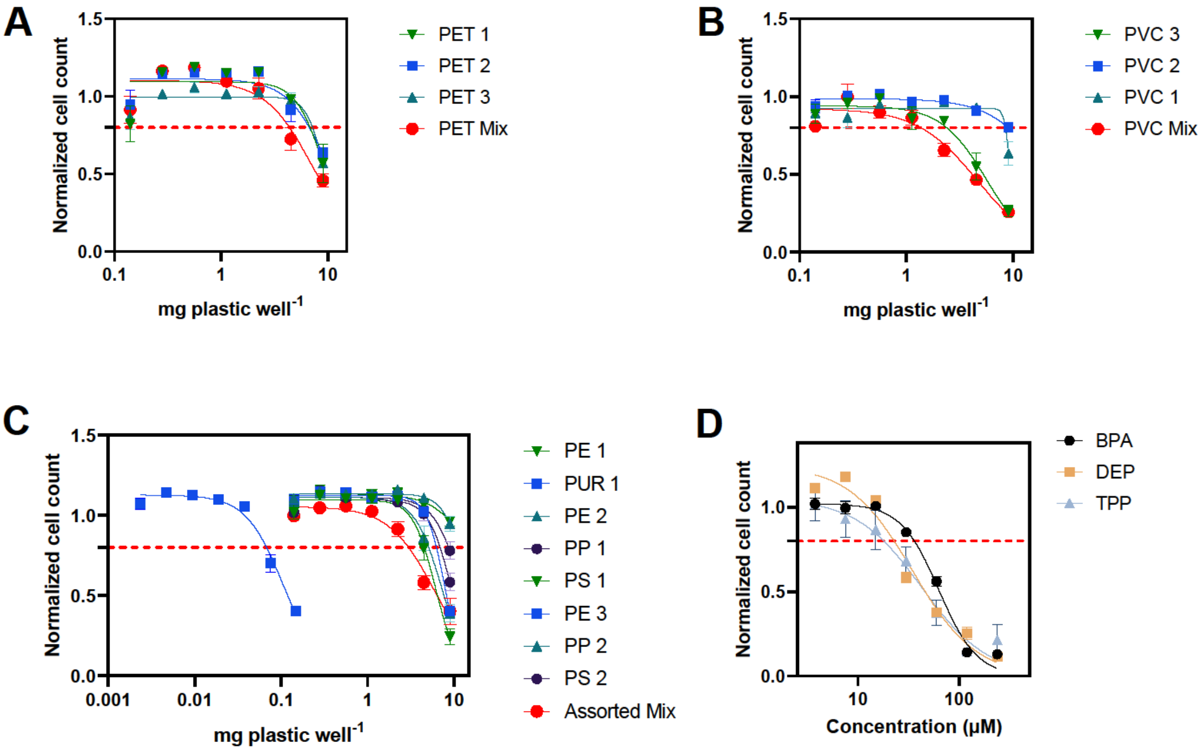


**Figure S6. Cytotoxicity of non-transfected HTLA cells exposed to (A) three PET extracts, and their PET Mix, (B) three PVC extracts and their PVC Mix, (C) eight extracts from PP, PE, PS, and PP and their Assorted Mix, and (D) plastic chemicals BPA (black), DEP (yellow), and TPP (grey).** The red horizontal line represents the cytotoxicity cut-off of 0.8. Concentration (x-axis) is given in plastic mg plastic well$^{-1}$ for A-C and in μM for D. Data is normalized to the cell count of controls and shown in mean cell count ± SEM of four technical replicates and three biological replicates.

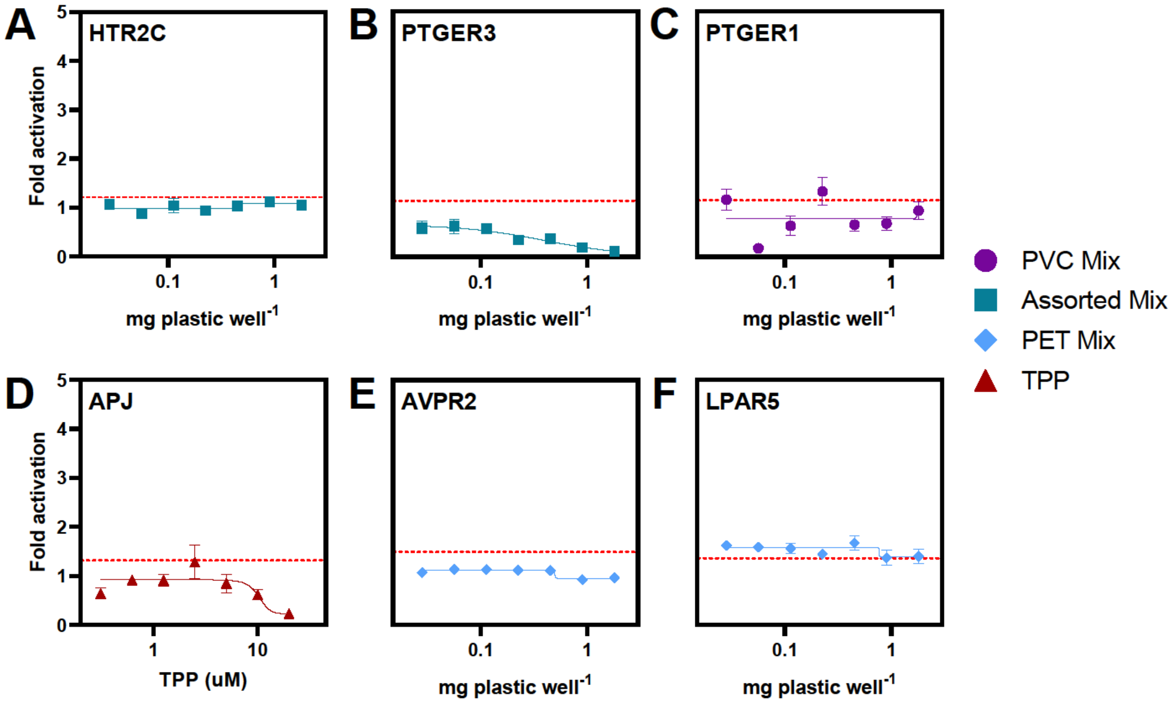


**Figure S7. The secondary PRESTO-Tango screen hits that could not be confirmed in a dose dependent manner.** The red horizontal line represents the LOD. Data are shown in mean fold activation ± SEM of three biological replicates with four technical replicates, each.

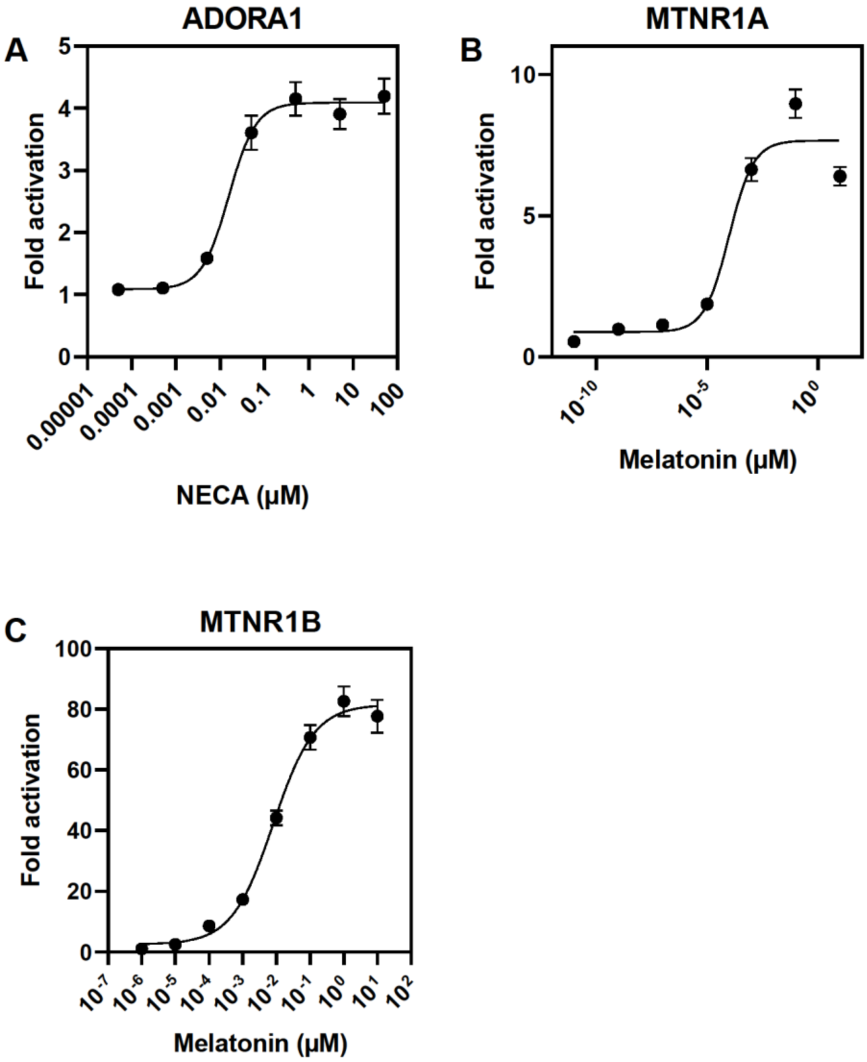


**Figure S8. Dose response relationship for (A) ADORA 1 exposed to reference compound NECA, (B) MTNR1A and (C) MTNR1B exposed to reference compound melatonin.** MTNR1B activation by melatonin served as the positive control for the primary and secondary PRESTO-Tango screens. 60 or more replicates per concentration (n ≥60). ADORA1 and MTNR1A activation by NECA and melatonin, respectively, served as the positive control for the analysis of the individual plastic extracts and the recovery assay. 48 or more replicates per concentration (n ≥48). Data are shown as mean ± SEM of fold activation calculated from CA of the SC and NC.

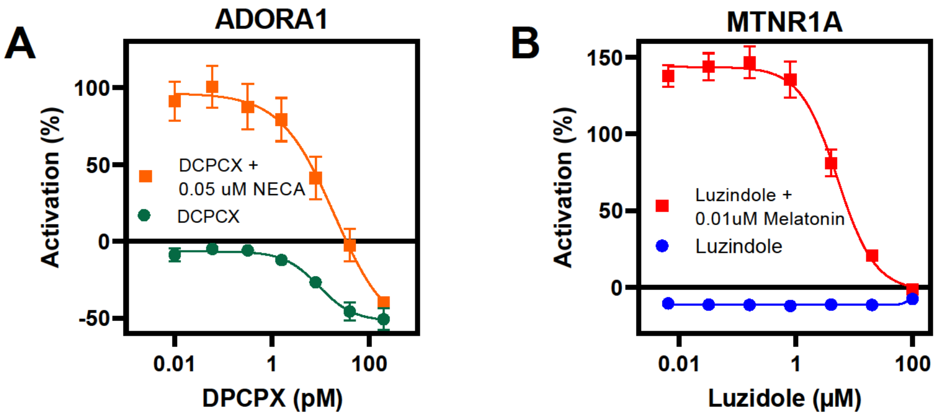


**Figure S9. Dose-response relationship for (A) ADORA1 and (B) MTNR1A exposed to reference antagonists DCPCX and luzindole.** ADORA1 was exposed to only DCPCX (green) and co-exposed with NECA (0.05 uM) and DCPCX (orange). Similarly, MTNR1A exposed to only luzidole (blue) was compared to co-expsoure with melatonin (0.01uM) and luzidole**.** ADORA1 and MTNR1A deactivation by DCPCX and luzidole, respectively, served as the positive control for the pharmacological knockdown confirmation. 12 or more replicates per concentration (n ≥12). Data are shown as mean ± SEM of activation normalized to the positive control (ADORA1 activation by NECA, MTNR1A activation by melatonin) and the CA (calculated as the mean of the SC and NC).

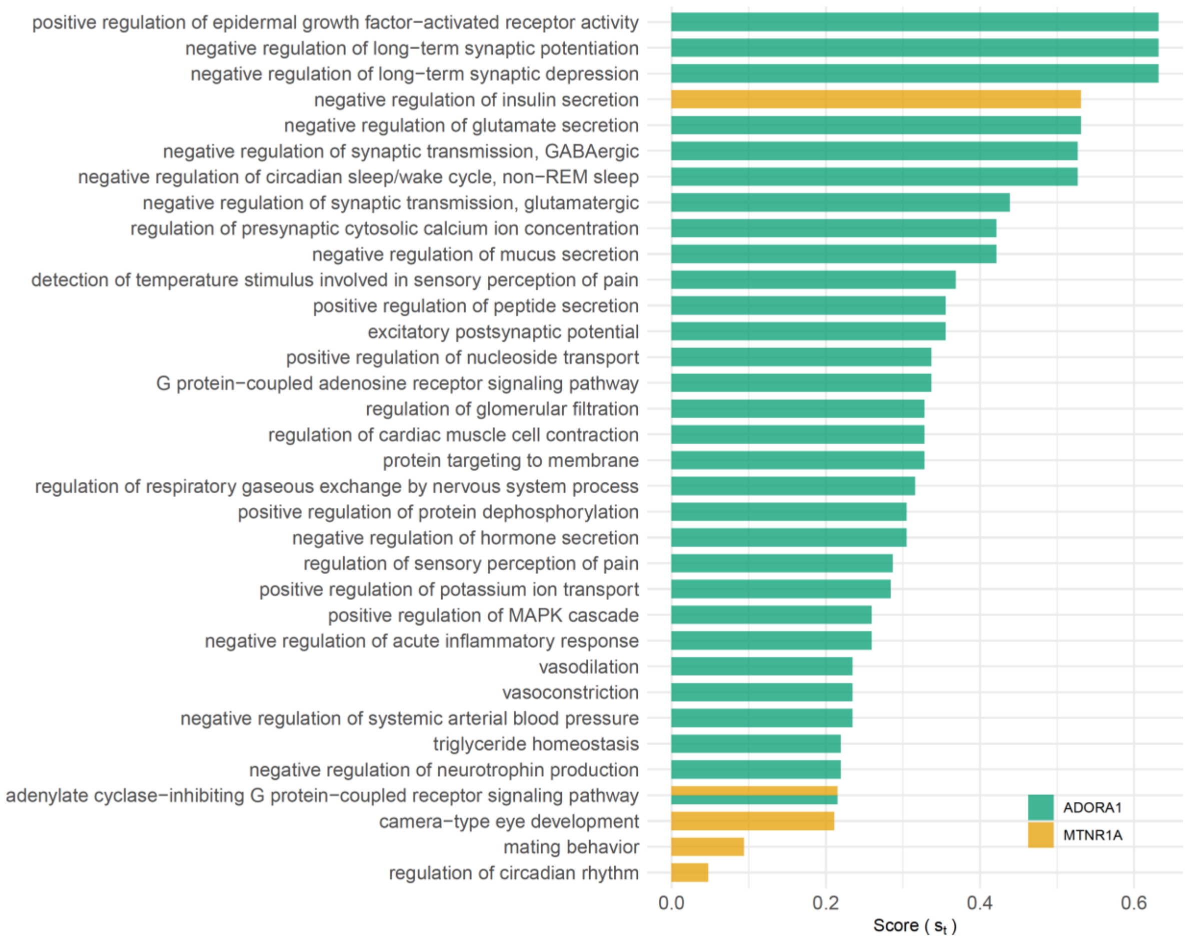


**Figure S10. Biological processes regulated by ADORA1 and MTNR1A**. GO terms of the 30 top ranked annotations for ADORA1 (green) and 5 annotations for MTNR1A (yellow). Each bar represents the structure-based ranking score ($S_t$) that indicates the biological specificity of a GO term.
